# Characterisation of virulence of *Puccinia coronata* f. sp. *avenae* in Australia in the 2023 growing season

**DOI:** 10.1101/2024.09.25.614061

**Authors:** Duong T. Nguyen, Eva C. Henningsen, David Lewis, Rohit Mago, Jana Sperschneider, Eric Stone, Peter N. Dodds, Melania Figueroa

## Abstract

Crown rust caused by the basidiomycete fungus *Puccinia coronata* f. sp. *avenae* (*Pca*) results in significant crop losses worldwide. Genetic solutions to protect against this disease require disease resistance gene discovery and introduction of resistance genes into elite germplasm by breeders. To inform disease resistance breeding activities, it is paramount to monitor changes of virulence in the *Pca* population and link those to genotypes of the pathogen. In 2023, a collection of 37 *Pca* isolates from diverse regions in Australia were gathered and their infection types across a commonly used set of oat differential lines determined to assign virulence pathotypes. We compared those virulence phenotypes to data collected in previous collections. While some of the virulence phenotypes had been reported in 2022, our analysis detected new virulence or an increase in frequency for virulence on some resistance genes. Notably, some of the frequency increases in virulence were recorded in Western Australia (WA), a region of interest due to its role in oat production for milling.

## Manuscript body

Oat crown rust, caused by *Puccinia coronata* f. sp. *avenae* (*Pca*), is a major foliar disease that is linked to substantial decreases in yield and grain quality of oats around the world, including Australia (GRDC 2016, Nazareno et al. 2018). Characterisation of the virulence of *Pca* populations is important to develop effective disease management strategies and inform breeding activities. For instance, pathogen monitoring can inform whether resistance genes in oat cultivars will remain effective in providing protection against the most prevalent genotypes of the pathogen and influence decisions such as which oat cultivars should be deployed or removed from fields due to disease susceptibility.

As in other biotrophic pathosystems, the interaction of oat and *Pca* results in an ‘arms-race’ scenario that is controlled by the gene-for-gene relationship between host resistance (*R*) genes and pathogen avirulence (*Avr*) genes (Dodds 2023; Flor 1971, Kaur and Bhatia 2021; Thrall et al. 2012). Under these dynamics, the presence of an *R* gene in the plant exerts pressure on the pathogen to evolve a variant (allele) of the *Avr* gene to enable plant infection and colonisation. The molecular and biochemical basis of disease resistance to rust fungi is the direct or indirect recognition interaction between an immunoreceptor, encoded by the *R* gene in the plant, and an *Avr* gene product (known as *Avr* effector) secreted by the pathogen. In the absence of such recognition, disease develops because the signalling cascade that results in immunity is not activated. In oat, close to 100 *R* genes to protect against crown rust disease have been catalogued. However, only a few of these *R* genes have been mapped to oat chromosomes and none has been cloned (Admassu-Yimer et al. 2018, 2022; Bush and Wise 1998; Chen et al. 2006; Gnanesh et al. 2013, 2015; Hoffman et al. 2006; Kebede et al. 2019; Klos et al. 2017; Kulcheski et al. 2010; McCartney et al. 2011; McNish et al. 2020; Satheeskumar et al. 2011; Sowa and Paczos-Grzeda 2020; Wight et al. 2005; Zhao et al. 2020). Similarly, the genomic location of multiple *Avr* effectors of *Pca* has been predicted through association mapping but no *Avr* genes have been positively identified yet (Hewitt et al. 2023; Miller et al. 2020).

The analysis of disease reactions across a set of oat genotypes (lines) containing different *R* genes after *Pca* inoculation is a common practice to assign race pathotypes and provide insights into the population structure and diversity of the pathogen (Carson 2011; Chong et al. 2000). One of the key success factors in race assignment (also known as pathotyping) is the access to host lines which carry distinct *R* genes. Unfortunately, the oat differential panels for race assignment in *Pca* still require molecular characterisation. Only a few near-isogenic lines have been developed making it difficult to ascertain if other *R* genes are present in the lines and could interfere with disease scoring (Hewitt et al. 2023; Nazareno et al. 2018; Nguyen et al. 2023). Nevertheless, a pathotyping approach can be valuable when coupled with genotypic information of the pathogen. There are limitations faced by only considering races as indicators of genetic diversity. These include the masking effects of dominant traits like avirulence in an organism that contains two nuclei and complex genetic interactions such as suppression of phenotypes that can manifest in both the plant and the pathogen (Figueroa et al. 2020; Jones and Dangl 2006; Petit-Houdenot and Fudal 2017).

Multiple fully phased nuclear haplotype genome assemblies of *Pca* isolates have recently become available that capture the chromosome composition of both haploid nuclei (Henningsen et al. 2022, 2024a). Furthermore, numerous genotypic lineages of the pathogen have been defined for *Pca* populations in Australia, Taiwan, USA, and South Africa (Henningsen et al. 2024a; Hewitt et al. 2023; Miller et al. 2020). Analysis of *Pca* isolates collected from 2020 to 2023 in Australia defined 18 unique lineages (Henningsen et al. 2024a). Importantly, it has been noted that similarities in virulence phenotypes do not necessarily reflect genotypic lineages, indicating the value of using both approaches to fully understand the population dynamics of *Pca*.

To expand the collection of *Pca* isolates reported in 2022 (Henningsen et al. 2024b), we connected with plant pathologists and industry representatives and obtained 37 *Pca*-infected leaf samples in 2023 from various oat-growing regions across Australia: 9 from Western Australia (WA), 10 from New South Wales (NSW), 8 from Victoria (VIC), and 10 from South Australia (SA). The isolates were revived by inoculation onto two broadly susceptible oat cultivars (‘Swan’ and ‘Marvelous’) following the methods outlined by Miller et al. (2020) and Henningsen et al. (2024b).

To assign virulence pathotypes to these of *Pca* isolates, we utilised the oat differential sets recently compiled by Henningsen et al. (2024b) with lines sourced from USDA-ARS (St. Paul, MN, USA) and the Australian Grain Genebank (AGG). We used a total of 49 oat lines (Supplementary **Table S1**), after excluding those that were no longer effective against most *Pca* isolates in Australia (Henningsen et al. 2024b) and those whose genetic discrepancies suggest potential seed stock differences or *R* gene segregation (Nguyen et al. 2023). This report adds information to data shared by Henningsen et al. (2024a), which used 27 Australian differential lines to establish virulence profiles for 19 out of these 37 *Pca* isolates from 2023. Infection and scoring methods were previously described by Miller et al. (2020), Nazareno et al. (2018), and (Henningsen et al. 2024b). The raw data were converted to a 0 to 9 numeric scale (Supplementary **Table S2**) to create heatmaps for visualisation in R (4.2.2) using the package “ComplexHeatmap” (Gu et al. 2016).

Using a 10-letter race coding system as previously described by Carson (2011), Chong et al. (2000), and Nazareno et al. (2018), we detected 32 unique races among the 37 *Pca* isolates (**Table 1**), confirming previous findings of high diversity in virulence phenotypes in Australia (Henningsen et al. 2024b). Overall, the most broadly virulent isolates in 2023 were from the state of VIC (23VIC09 – race TQMPNBTPM-) followed by isolates from WA (23WA07 and 23WA43 – race TQRPLBTKL-, and 23WA20-race TQTPLBTKL-) and SA (23SA41, 23SA42 – race TQRPLBTKL-), which were virulent to 31 and 28 differential lines, respectively (**Table 1**). The least virulent isolates were from NSW (23NSW13 – race BBBBBBDCB- and 23NSW36 – race GDBBBBNCB-), which were virulent to only two and five differential lines, respectively.

**Table 1.**
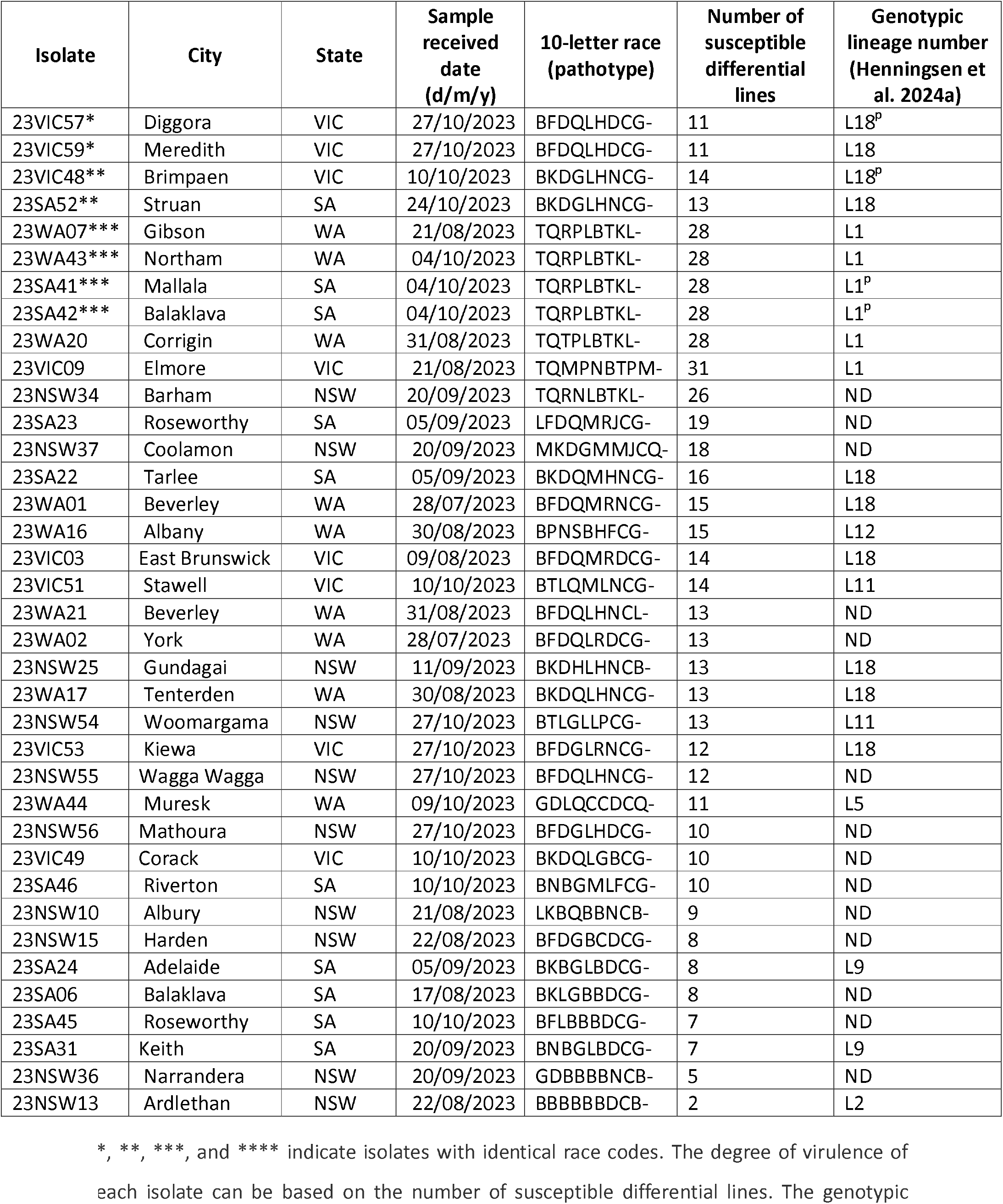
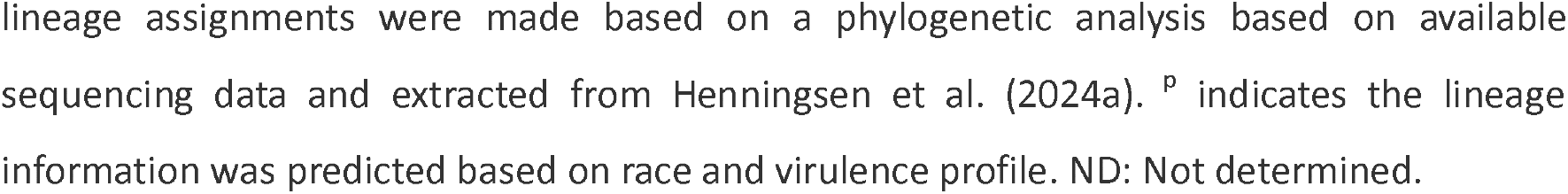
List of *Puccinia coronata* f. sp. *avenae* (*Pca*) isolates collected across Australia 2023.

Two representative *Pca* isolates were found for the races, “BFDQLHDCG-”, “BKDGLHNCG-”, while four isolates of race “TQRPLBTKL-” were detected. Notably, one of these races (“BKDGLHNCG-”) was found across different regions (SA and VIC, see isolates 23SA52 and 23VIC48). Similarly, four isolates from SA (23SA41 and 23SA42) and WA (23WA07 and 23WA43) all exhibited the race “TQRPLBTKL-” (**Table 1**). This wide geographic distribution of races was also noted in 2022 when several other isolates with identical races were found between NSW and VIC (BFDGLHBCQ-), as well as NSW and QLD (GBBQBBLFL-) (Henningsen et al. 2024b). We did not detect any races in common between isolates collected in 2022 (Henningsen et al. 2024b) and 2023. However, most isolates within each year also had unique phenotypes (2022-42 of 48; 2023-32 of 37), suggesting that neither collection is fully sampling the phenotypic diversity of the pathogen in Australia. Thus, it is difficult to distinguish whether the year-to-year differences reflect a change in the *Pca* population or result from insufficient sampling depth. A larger sample size may be necessary to accurately determine the frequency of virulence traits.

In 2023, five oat lines, Barcoo, Saia, WIX4361.9, Pc59, and Pc63, were resistant to all *Pca* isolates across the country (**Fig. 1**). Among these, only Barcoo and Pc63, also displayed resistance across the country in 2022 (Henningsen et al. 2024b). However, we note that virulence to Pc63 and Barcoo was previously documented in 2001 (Brake et al. 2001) and 2010 (Park and Wellings 2010), suggesting that virulence frequencies to *R* genes present in these two lines remain rare and sampling biases may be playing a role in the lack of detection of the virulence. The oat cultivar Barcoo was postulated to carry genes *Pc39, Pc61*, and *PcBett* (Park 2013). Nonetheless, our findings suggested the resistance of Barcoo was due to the presence of other or additional gene(s), as we found that differential lines Pc39, Pc61, and Bettong (*PcBett*) were all broadly susceptible to *Pca* isolates from 2023.

**Fig. 1.**
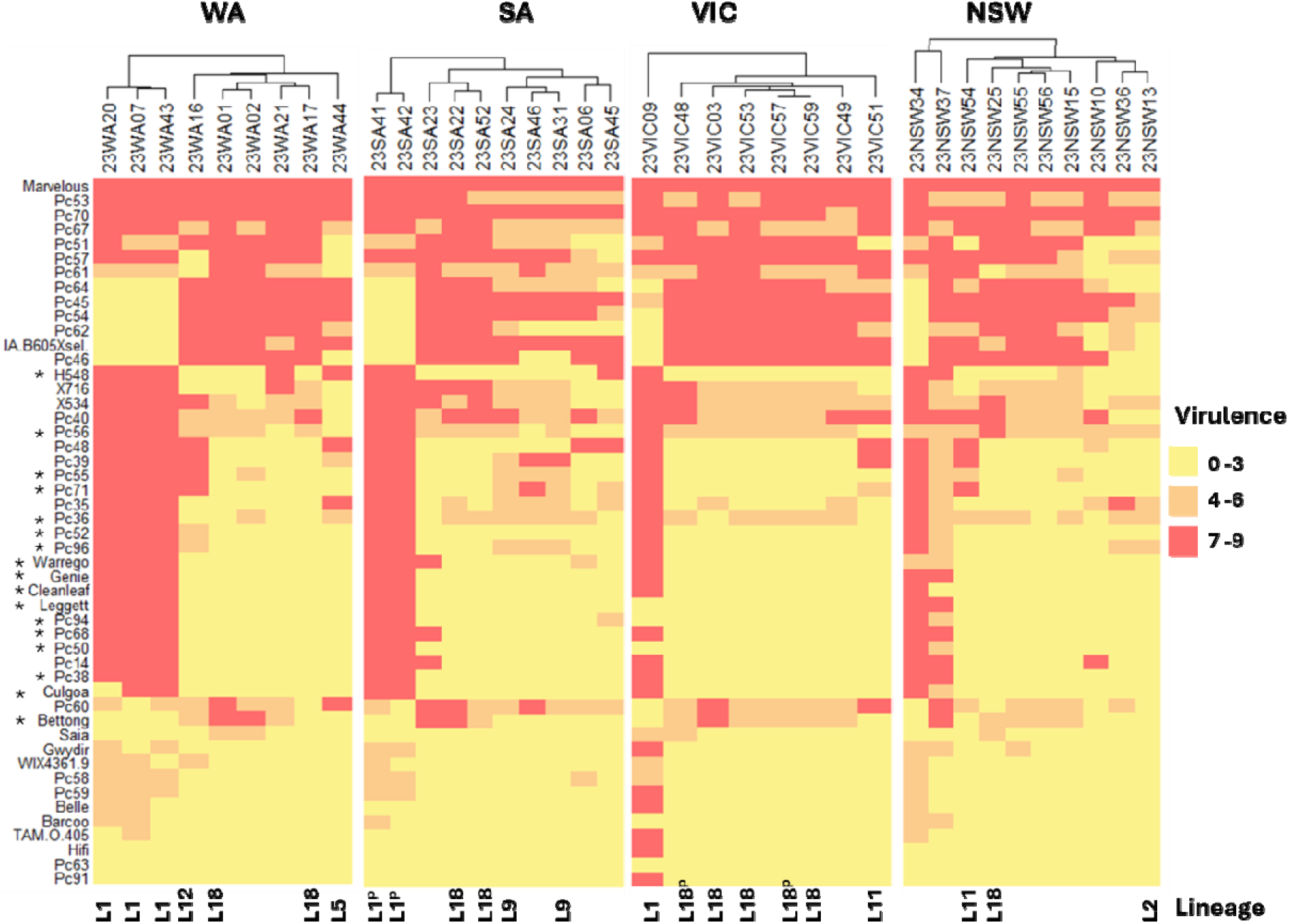
Heatmap showing virulence profiles of *Pca* isolates collected in 2023 (x-axis) on the oat differential lines (y-axis). High infection scores indicating high virulence (susceptibility) are shown in red, and lower infection scores indicating avirulence (resistance) are shown in yellow and orange. Columns are ordered by hierarchical clustering divided by regions. * show oat lines resistant to WA *Pca* isolates in 2022 (Henningsen et al. 2024b) but susceptible in 2023. Lineage (L) numbers of *Pca* isolates were extracted from a phylogenetic analysis reported by Henningsen et al. (2024a). ^p^ indicates the lineage information was predicted based on virulence profile, requiring further confirmation.

Eleven oat lines showed disease resistance to all WA isolates in 2023 (**Fig. 1**). Among these, eight lines including Barcoo, Gwydir, HiFi, Saia, WIX 4361-9, Pc59, Pc63, and Pc91 aligned with data released by the Australian Cereal Rust Control Program (ACRCP), that reported no virulence to these lines in WA isolates (Australian Cereal Rust Survey, https://www.sydney.edu.au/science/our-research/research-areas/life-and-environmental-sciences/cereal-rust-research/rust-reports.html). However, virulence to these lines have already been detected in the eastern parts of the country previously (Australian Cereal Rust Survey 2020, 2021, 2022; Brake et al. 2001; Henningsen et al. 2024b, Park and Wellings 2010). This poses a risk due to the potential migration of rust spores which can be easily spread by air currents (Simon 1985).

The *Pca* isolates collected from WA in 2023 showed increased virulence compared to isolates from 2022 (Henningsen et al. 2024b). While 27 oat differential lines were classified to be resistant to all isolates of *Pca* from WA in 2022, 17 of these oat lines were susceptible to WA isolates in 2023 (**Fig. 1**). Among them, nine lines Bettong, Cleanleaf, Culgoa, Genie, Warrego, Pc36, Pc50, Pc56, and Pc68 were identified as susceptible to WA isolates for the first time, with previous ACRCP surveys reporting no virulence to these lines in WA (Australian Cereal Rust Survey). The observation that more differential lines were susceptible to WA isolates in 2023, despite phenotyping fewer isolates in 2023 (n=9) compared to 2022 (n=20) (Henningsen et al. 2024b), suggests a shift in the population of *Pca* in WA. Such an increase in virulence could be explained by the evolution of the *Pca* isolates or the migration of isolates from other states to WA. However, it is also possible that we missed detecting these virulence traits in 2022.

We recently included 19 *Pca* isolates from the 2023 collection in a study using genome-wide sequencing data to determine the phylogenetic relationships and genotypic diversity within the Australian *Pca* population and compare them to other parts of the world (Henningsen et al. 2024a). The study reported the presence of 18 *Pca* genotypic lineages in Australia, with lineage L1 and L18 being the most prevalent based on the total number of their corresponding isolates across the country (Henningsen et al. 2024a). The similarities in virulence profile of the 23SA41 and 23SA42 to the lineage 1 isolates 23WA07 and 23WA43 suggest that they may also belong to this lineage L1 (**Table 1, Fig. 1**), while the isolates 23VIC57 and 23VIC48 may belong to lineage L18 as they display similar virulence profiles to isolates 23VIC59 and 23SA52, respectively (**Table 1** and **Fig. 1**). However, genome-wide sequencing data would be needed to confirm these predictions as we note that race information does not always correlate with lineage (Henningsen et al. 2024a).

Among the 2023-*Pca* population, lineage L18 contains the most races (n=7), while lineage L1 follows with three races (**Table 1**). Both lineages L1 and L18 were the most widespread in 2023, as they were represented in the western and eastern regions of the country (**Fig. 1**). According to the 2023 data, lineage L1 is highlighted as the most broadly virulent lineage in Australia. For example, isolate 23VIC09 (race TQMPNBTPM-) was virulent against 31 differential lines (**Table 1** and **Fig. 1**). These observations also agreed with the finding by Henningsen et al. (2024a) when analysing a larger collection of 137 isolates from 2020 to 2023.

In 2023, the detection of three lineage L1 isolates in WA (23WA07, 23WA20, 23WA43) may have contributed to the increased susceptibility of differential lines to *Pca* in this region (**Fig. 1**) compared to 2022 when no isolates of this lineage were detected (Henningsen et al. 2024a). A single *Pca* isolate belonging to Lineage L1 was identified in WA in 2020, but this isolate exhibited lower virulence levels compared to 23WA07, 23WA20, and 23WA43 based on phenotypic data on 27 Australian differential lines (Henningsen et al. 2024a). It may be the case that the highly virulent L1 isolates detected in WA in 2023 migrated from other regions since similar races were found in SA (isolates 23SA41 and 23SA42) (**Table 1** and **Fig .1**). Alternatively, the local L1 lineage may have mutated to acquire additional virulence traits in WA. These alternatives could be distinguished by phylogenomic analysis of these isolates.

Recent comparisons of Pca populations from Australia, the USA, and Taiwan revealed that the genotypic and phenotypic diversity of the Pca population in Australia is not as high as that in the USA, but is greater than that observed in Taiwan (Henningsen et al. 2024b; Hewitt et al. 2023; Ho et al. 2024; Miller et al. 2020). This diversity level is high enough to question if only somatic mutations and migration are contributing to the evolution of the pathogen (Figueroa et al. 2020). In the USA, the reassortment of virulence genes is widely facilitated by sexual reproduction and somatic hybridisation (Henningsen et al. 2024a; Hewitt et al. 2023; Miller et al. 2020). Sexual reproduction is enabled by the presence of an alternate host, such as buckthorn (Rhamnus cathartica) particularly in the North of the USA (Nazareno et al. 2018). While R. cathartica has not been found in Australia, other related Rhamnus species (e.g., R. alaternus) whose presence is reported in Australia (Stajsic 2023), can also serve as sexual hosts (Dinoor 1962). Through analysis of recombination signatures in the genomes of various Pca isolates (L9 – 23SA24, 23SA31; L11 – 23VIC51, 23NSW54; L18 – 23WA01, 23VIC03, 23WA17, 23SA22, 23NSW25, 23SA52, 23VIC53, 23VIC59), Henningsen et al. (2024a) provided support to the hypothesis that although rare, genetic recombination may occur in Australia. In Australia, the widespread distribution of wild oats significantly influences the epidemiology of oat crown rust (Chambers et al. 2022). Wild oats can become infected by crown rust, providing additional opportunities for mutation and adaptation (Chambers et al. 2022). Thus, it is imperative to employ molecular monitoring of Pca and investigate the population near clusters of Rhamnus species.

In conclusion, the ongoing expansion of the Pca collection is necessary to improve our understanding of phenotypic variations in Australia. This, in turn, will aid in the development of more effective plant protection strategies and the identification of new genetic resources for controlling oat crown rust.

## Acknowledgment

We thank Allan Rattey (InterGrain), Andrea Hills (DPIRD WA^1^), Daniel Malecki-Lee (DPIRD WA), Deb Donovan (DPIRD WA), Tara Garrard (SARDI^2^), and Peter Dracatos (La Trobe University) and their organizations for contributing samples to the collection used in this study.

^1^Department of Primary Industries and Regional Development, WA

^2^South Australian Research and Development Institute

